# *VviWRKY10* and *VviWRKY30* co-regulate powdery mildew resistance through modulating SA and ET-based defenses in grapevine

**DOI:** 10.1101/2023.12.08.570812

**Authors:** Min Zhou, Hongyan Wang, Xuena Yu, Kaicheng Cui, Yang Hu, Shunyuan Xiao, Ying-Qiang Wen

**Author notes:** Corresponding author (Y.-Q. Wen). These two authors contributed equally (Min Zhou, Hongyan Wang). The authors responsible for distribution of materials integral to the findings presented in this article in accordance with the policy described in the Instructions for Authors (https://academic.oup.com/plcell/pages/General-Instructions) is: Ying-Qiang Wen.

## Abstract

Grapevine is one of the most important economic fruit crops in the world. The widely cultivated grapevine (*Vitis vinifera*, *Vvi*) is susceptible to powdery mildew caused by *Erysiphe necator*. In this study, CRISPR-Cas9 was used to simultaneously knock out *VviWRKY10* and *VviWRKY30* encoding two transcription factors reported to be implicated in defense regulation. Fifty-three *wrky10* (*Vviwrky10* single mutant) transgenic plants and fifteen *wrky10wrky30* (*Vviwrky10Vviwrky30* double mutant) transgenic plants were generated. In a 2-year field evaluation of resistance to powdery mildew, the *wrky10* showed strong resistance, while the *wrky10wrky30* showed moderate resistance. Further analyses revealed that salicylic acid (SA) and reactive oxygen species (ROS) contents in the leaves of *wrky10* and *wrky10wrky30* were significantly increased, so did the ethylene (ET) content in the leaves of *wrky10*. The results from dual luciferase (DLUC) reporter assays, electrophoretic mobility shift assays (EMSA) and chromatin immunoprecipitation (ChIP)-qPCR assays demonstrated that VviWRKY10 could directly bind to the W-boxes in the promoter of *VviEDS5-2*, *VviPR1*, *VviPR5* and *VviRBOHD2* and inhibit their transcription, supporting its role as a negative regulator of SA-dependent defense. By contrast, VviWRKY30 could directly bind to the W-boxes in the promoter of *VviACS3* and *VviACS3L* and promote their transcription, playing a positive role in ET production and ET-dependent defense. Moreover, VviWRKY10 and VviWRKY30 can bind to each other’s promoters and mutually inhibit each other’s transcription. Taken together, our results have revealed a complex mechanism of regulation by VviWRKY10 and VviWRKY30 for activation of measured and balanced defense responses against powdery mildew in grapevine.

**One-sentence summary:** *VviWRKY10* and *VviWRKY30* play different roles in powdery mildew resistance through the salicylic acid and ethylene, respectively, and there is mutual inhibition between *VviWRKY10* and *VviWRKY30*.

## Introduction

Grapevine is an important fruit crop with high nutritional and economic values and a long history of cultivation in the world (Jaillon et al., 2007). However, the widely cultivated European grapevine species *Vitis vinifera* is highly susceptible to powdery mildew caused by *Erysiphe necator*, resulting in serious losses in growth and yield (Gadoury et al., 2012, Gao et al., 2016, Hu et al., 2018). Therefore, it is necessary and urgent to create disease-resistant grapevine through breeding and novel transgene technologies.

Rapid and massive transcriptional reprogramming events are often associated with plant-pathogen interactions, in which transcription factors (TFs) play important roles (Tsuda & Somssich, 2015). Among the large family of transcription factors in plants, WRKY TFs are mainly involved in responding to various biological stresses (Eulgem & Somssich, 2007, Jiang et al., 2017). In Arabidopsis, AtWRKY18 and AtWRKY40 are negative regulators of powdery mildew resistance by inhibiting the expression of *CYP71A13*, *EDS1* and *PAD4*. The double mutant *wrky18wrky40* and triple mutant *wrky18wrky40wrky60*, but not the *wrky18*, *wrky40*, and *wrky60* single mutants, are significantly more resistant to powdery mildew than the wild type (WT), indicating that AtWRKY18 and AtWRKY40 have partially redundant functions in Arabidopsis (Xu et al., 2006, Shen et al., 2007, Pandey et al., 2010). The *Xanthomonas* type-III effector XopS interacts with and inhibits proteasomal degradation of CaWRKY40a, preventing the plant from properly activating defense genes and the expression of key genes required for stomatal closure, making it easier for pathogens to enter the pepper leaves (Raffeiner et al., 2022). RhWRKY13, which is an ortholog of AtWRKY40, protects rose petals against *Botrytis cinerea* infection by binding to the promoter region of *RhCKX3* and *RhABI4*, inhibiting their expression in rose petals, increasing CK content and reducing ABA response (Liu et al., 2022).

Further, it was reported that the target genes of AtWRKY18 and AtWRKY40 under flg22 treatment included multiple hormone pathways, including SA and ET (Birkenbihl et al., 2017). It is well known that WRKYs can regulate hormone pathways and play important roles in plant immunity. CaWRKY27 enhances the resistance of tobacco to *Ralstonia solanacearum* by regulating SA, JA and ET-mediated signaling pathways (Dang et al., 2014). AtWRKY55 regulates leaf senescence and pathogen resistance by directly binding to the W-boxes of *RBOHD*, *ICS1*, *PBS3*, and *SAG13*, activating their expression and integrating ROS and SA pathways in Arabidopsis (Wang et al., 2020).

It was reported that *WRKY10* and *-30* in grapevine, as homologous genes of *AtWRKY18* and *-40*, also are involved in regulating the interaction between grapevine and pathogens (Guo et al., 2014, Ma et al., 2021). Overexpression of *Vitis amurensis WRKY10* in *Arabidopsis thaliana* and in *V. vinifera* increases resistance to *Botrytis cinerea* (Wan et al., 2021). Due to different naming rules, some studies refer to *VviWRKY30* as *VviWRKY40*. *Plasmopara viticola* effector PvRXLR111 interacts with VviWRKY40 and improves its stability to suppresses the immune response of grapevine to *Plasmopara viticola* (Ma et al., 2021). Besides the above studies, it remains to be seen whether *VviWRKY10* and *-30* regulate resistance to powdery mildew in grapevine, and whether they also have partial functional redundancy as do *AtWRKY18* and *-40* in Arabidopsis.

In this study, two mutants in grapevine, *wrky10* and *wrky10wrky30*, were obtained and used for characterizing their potential role in resistance to powdery mildew. It was found that *wrky10* showed strong resistance and *wrky10wrky30* showed moderate resistance. Moreover, the resistance of the mutants was due to significant increase of SA and ROS levels in *wrky10* and *wrky10wrky30* compared to wild-type plants. Interestingly, *wrky10* but not *wrky10wrky30* exhibited high-level ET accumulation. By DLUC, EMSA and ChIP-qPCR, we proved that VviWRKY10 inhibits *VviEDS5-2*, *VviPR1*, *VviPR5* and *VviRBOHD2* expression via directly targeting their promoters. We also showed that VviWRKY30 binds to the promoters of *VviACS3* and *VviACS3L* and promote their expression, and VviWRKY10 and VviWRKY30 mutually inhibit each other’s gene expression. Combined, our results demonstrate distinctive roles of *VviWRKY10* and *VviWRKY30* in modulating grapevine’s defense responses against powdery mildew through transcriptional regulation of different genes involved in the SA, ET and ROS pathways.

## Results

### Identification and characterization of *VviWRKY10* and *VviWRKY30* in grapevine

To investigate whether *VviWRKY10* and *VviWRKY30* are responsive to powdery mildew, cDNA of these two genes was prepared from leaf tissues of the susceptible *V. vinifera* cv. Cabernet Sauvignon. Analysis of the deduced protein sequences showed that both VviWRKY10 and VviWRKY30 had a highly conserved WRKY domain and a typical C_2_H_2_ zinc finger motif (Supplemental Figure S1). The phylogenetic tree was constructed by using the protein sequences of VviWRKY10, VviWRKY30 and WRKYs related to Arabidopsis, apple, pepper and rice. The results showed that VviWRKY30 was closely related to MdWRKY40, clustered with CaWRKY40, AtWRKY40, and VviWRKY10 into one subfamily (Figure 1A). Gene expression of *VviWRKY10* and *VviWRKY30* in ‘Cabernet Sauvignon’ inoculated with powdery mildew *E. necator* NAFU1 (*En*. NAFU1) (Gao et al., 2016) was detected by quantitative real-time reverse transcription PCR (qRT-PCR). The transcript level of *VviWRKY10* had a rapid increase from 0 to 24 hours post inoculation (hpi), and peaked at 24 hpi, which was 8.7-fold than at 0 hpi (Figure 1B). Differing from *VviWRKY10*, expression of *VviWRKY30* increased from 24 to 48 hpi, peaking at 48 hpi with a 71-fold increase relative to its expression at 0 hpi (Figure 1C). Hence, in terms of gene transcription, *VviWRKY10* responded to powdery mildew infection earlier than *VviWRKY30*.

**Figure 1.**
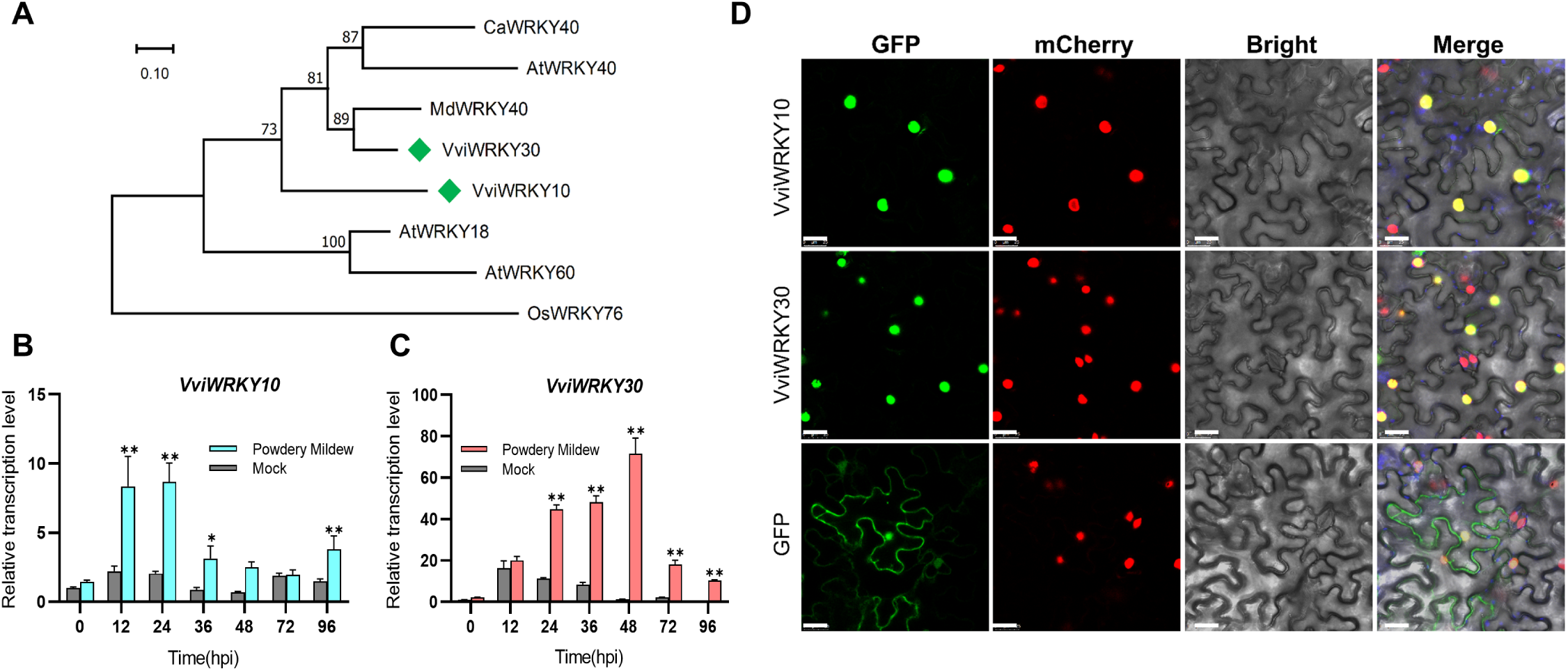
Phylogenetic and expression analysis of *VviWRKY10* and *VviWRKY30*. (**A**) Phylogenetic analysis of WRKYs proteins in *Vitis vinifera*(*Vvi*), *Arabidopsis thaliana* (*At*), *Oryza sativa*(*Os*), *Malus domestica*(*Md*), and *Capsicum annuum*(*Ca*). The green diamond squares represent the WRKYs proteins in *Vitis vinifera*. (**B**-**C**) Relative transcript levels of *VviWRKY10*(**B**) and *VviWRKY30*(**C**) post *Erysiphe necator* NAFU1 inoculation*. VviACTIN7* (XM_002282480.4) gene was used as an endogenous control. (**D**) Subcellular localization of VviWRKY10 and VviWRKY30. AtH2B:mCherry indicated the nuclear marker. Bars, 25 μm. Each data point represents the mean ± standard deviation of three biological replicates, and asterisks indicate significant difference compared with 0 h post inoculation (hpi) (Two-way ANOVA, **= P < 0.01; *= P < 0.05).

To determine if VviWRKY10 and VviWRKY30 have nuclear localization like most transcription factors, we constructed the *35S:VviWRKY10-eGFP* (enhanced Green Fluorescent Protein) and *35S:VviWRKY30-eGFP* fusion expression vectors and transiently expressed in leaves of *Nicotiana benthamiana*, using *35S:AtH2B-mCherry* as the nuclear localization marker. As shown in Figure 1D, VviWRKY10 and VviWRKY30 had typical nuclear localization, which was also verified by overlapping localization with the marker protein.

### CRISPR-Cas9-mediated mutagenesis of *VviWRKY10* and *VviWRKY30*

It was reported that the *Atwrky18/40* double and *Atwrky18/40/60* triple mutants but not the *Atwrky18*, *Atwrky40*, or *Atwrky60* single mutants were almost fully resistant to powdery mildew (Shen et al., 2007, Pandey et al., 2010). Therefore, we constructed a dual-target vector targeting *VviWRKY10* and *VviWRKY30* simultaneously in grapevine. The two target regions were located on the first exon of *VviWRKY10* and the third exon of *VviWRKY30*, respectively (Figure 2A). Two small guide (sg) RNAs were integrated into an intermediate vector to form an expression box, which was eventually reassembled into a GFP-tagged vector (Figure 2B). The vector was transformed into ‘Cabernet Sauvignon’ proembryo masses (PEM) by *Agrobacterium*-mediated transformation, and the transgene-positive plants were reported by GFP signals in the subculture (Supplemental Figure S2, A and B). The results indicate that GFP fluorescence remains stable in the tissue-cultured plantlets even after transplanting them into a greenhouse (Supplemental Figure S2, C and D).

**Figure 2.**
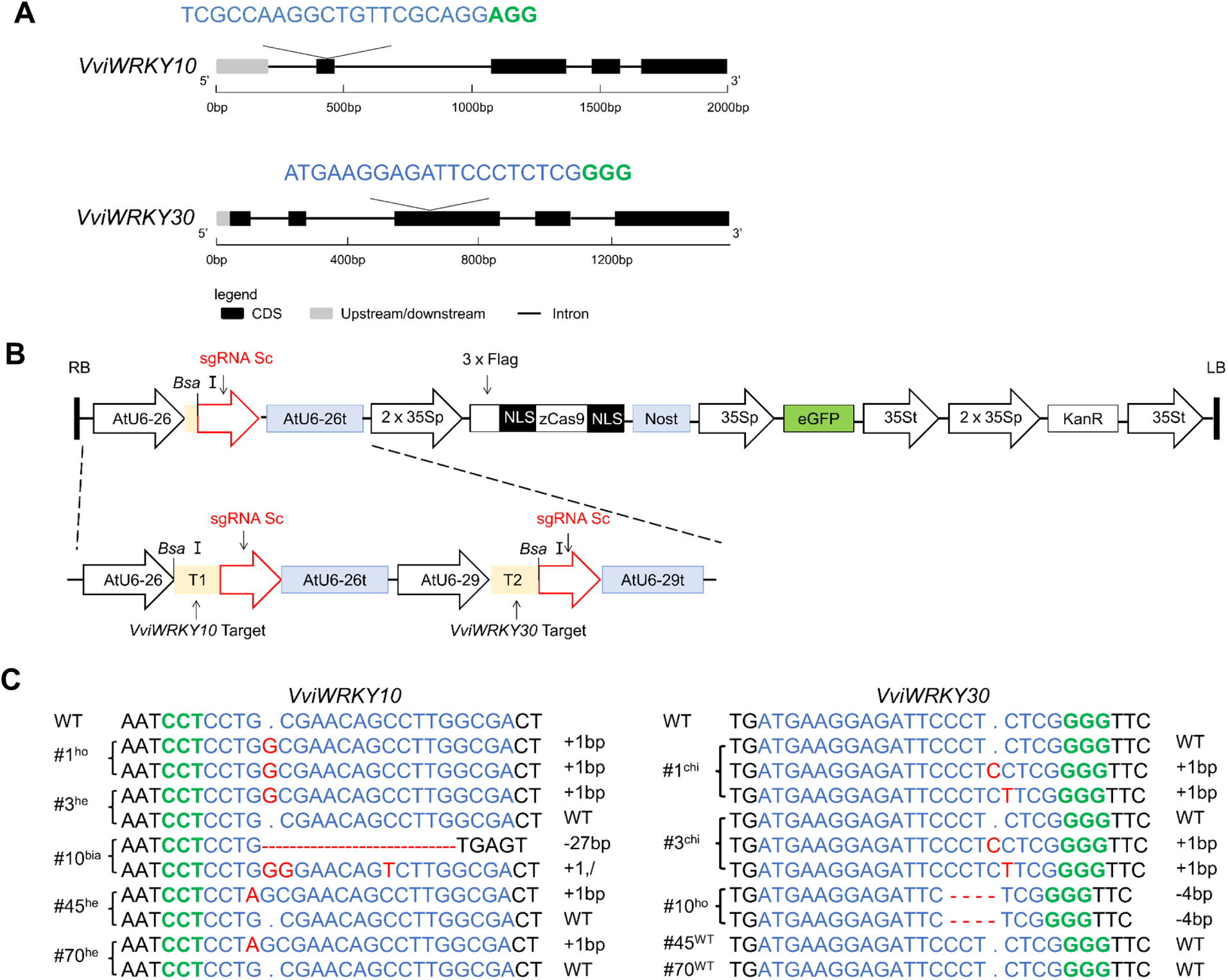
Construction of knockout vector and detection of edited plants. (**A**) The position of the target sites in *VviWRKY10* and *VviWRKY30*. (**B**) Schematic diagram of an expression cassette of intermediate vector and T-DNA region of pKSE401-GFP expression vector. (**C**) Different types of CRISPR/Cas9-induced mutations were detected in *VviWRKY10* and *VviWRKY30* genes of mutants. Type of mutations with “–” for deletion, “+” for insertion, “/” for substitution. ^ho^ Homozygous, ^he^ Heterozygous, ^bia^ Biallelic, ^chi^ Chimeric. The red, green and blue letters indicated mutation sequences, PAM sequences and target sequences respectively.

A total of 111 regenerated plantlets were obtained, of which 91 were GFP positive and the transformation efficiency was 81.98%. Sixty-eight mutants were obtained from 91 GFP-positive plants, including 53 *wrky10* mutants and 15 *wrky10wrky30* double mutants with an editing efficiency of 74.73% (Table 1, Supplemental Figure S3A). However, no *wrky30* mutants were obtained despite repeated transformation efforts. Among *wrky10* mutants, different types of mutations were identified. They included biallelic, heterozygous, and chimeric mutations as shown in lines 34, 45, 66, 70 and 13; Among the *wrky10wrky30* double mutants, there are several different types of combinatory mutations in the two target genes. For example, the mutations in line 1 is homozygous for *VviWRKY10* and chimeric for *VviWRKY30*; the mutations in line 10 and line 59 are biallelic for *VviWRKY10* and homozygous or chimeric for *VviWRKY30*; the mutations in line 3 and line 26 are the heterozygous for *VviWRKY10* and chimeric for *VviWRKY30* (Figure 2C, Figure 3A, Supplemental Figure S3B, Supplemental Figure S4, Table 1).

**Figure 3.**
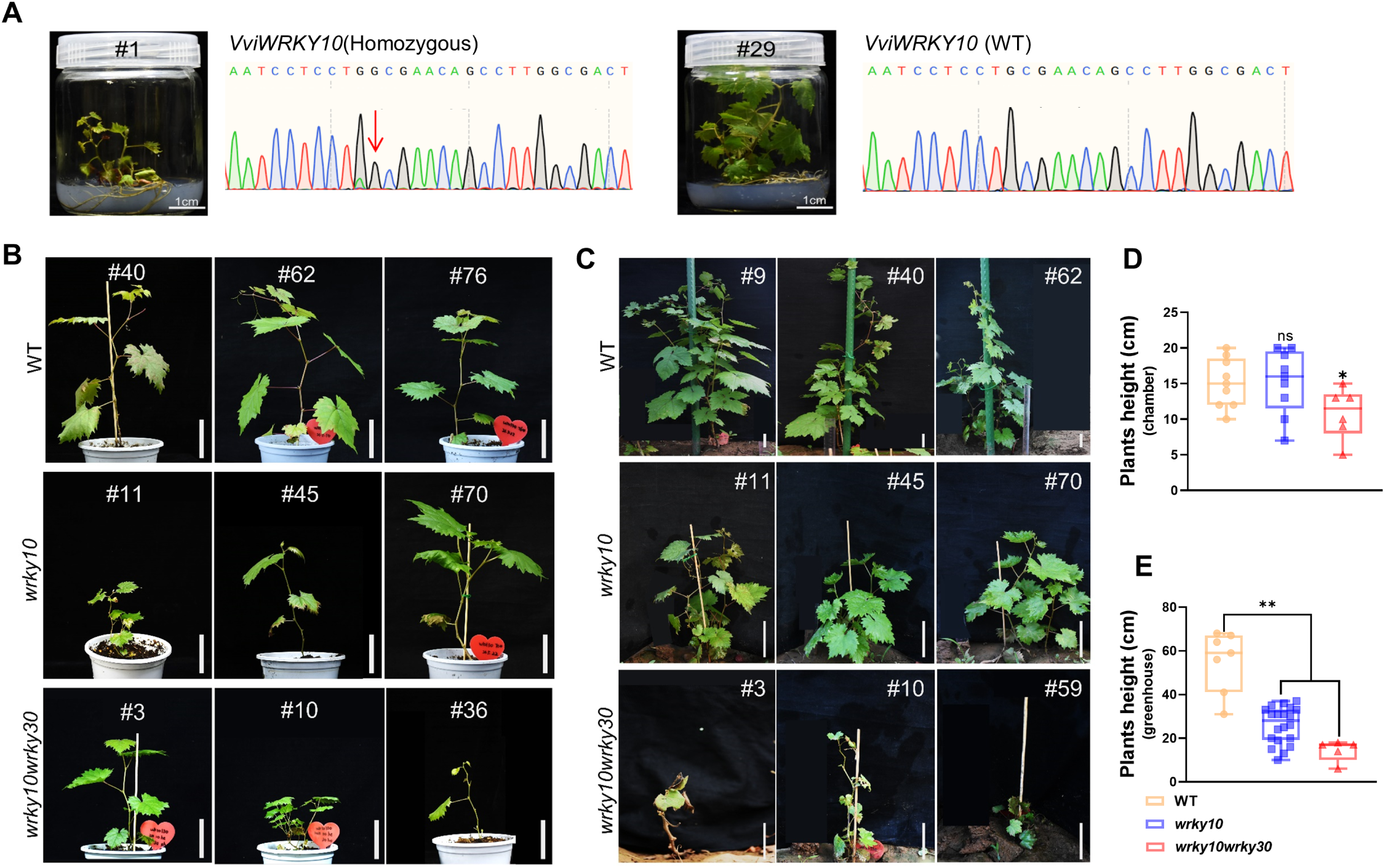
CRISPR-Cas9-mediated mutagenesis of *VviWRKY10* and *VviWRKY30* regulate growth in grapevine. (**A**) Phenotype and sequencing chromatograms of #1 and #29, insertion G was indicated by red arrow. (**B**-**C**) Phenotypes of WT and mutants grown in chamber(**B**) and greenhouse(**C**). Bars, 5 cm. (**D**-**E**) Plants height of WT and mutants grown in chamber(**D**) and greenhouse(**E**). Each data point represents the mean ± standard deviation of multiple biological replicates, and asterisks indicate significant difference compared with WT (Student’s *t* test, **= P < 0.01; *= P < 0.05; ns=no significant difference).

**Table 1.**
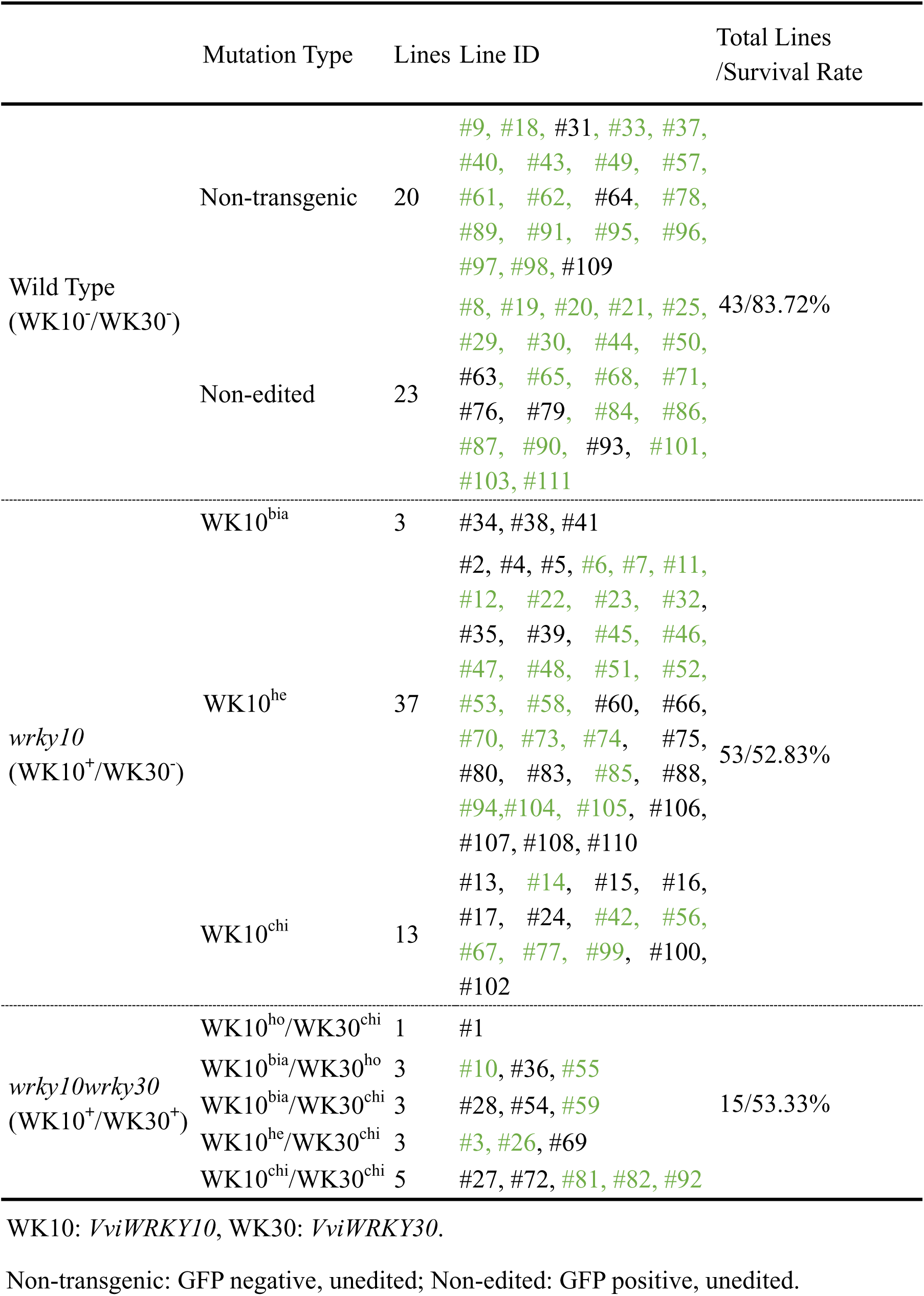
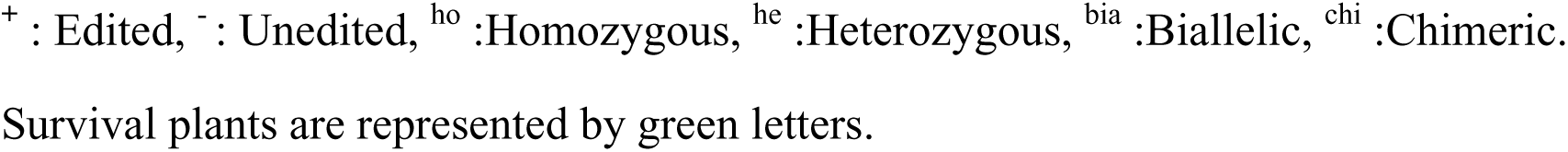
Summary of genome editing results in regenerated plants.

|  | Mutation Type | Lines | Line ID | Total Lines<br>/Survival Rate |
| --- | --- | --- | --- | --- |
| Wild Type<br>(WK10<sup>-</sup>/WK30<sup>-</sup>) | Non-transgenic | 20 | #9, #18, #31, #33, #37,<br>#40, #43, #49, #57,<br>#61, #62, #64, #78,<br>#89, #91, #95, #96,<br>#97, #98, #109 | 43/83.72% |
|  | Non-edited | 23 | #8, #19, #20, #21, #25,<br>#29, #30, #44, #50,<br>#63, #65, #68, #71,<br>#76, #79, #84, #86,<br>#87, #90, #93, #101,<br>#103, #111 |  |
| *wrky10*<br>(WK10<sup>+</sup>/WK30<sup>-</sup>) | WK10<sup>bia</sup> | 3 | #34, #38, #41 | 53/52.83% |
|  | WK10<sup>he</sup> | 37 | #2, #4, #5, #6, #7, #11,<br>#12, #22, #23, #32,<br>#35, #39, #45, #46,<br>#47, #48, #51, #52,<br>#53, #58, #60, #66,<br>#70, #73, #74, #75,<br>#80, #83, #85, #88,<br>#94, #104, #105, #106,<br>#107, #108, #110 |  |
|  | WK10<sup>chi</sup> | 13 | #13, #14, #15, #16,<br>#17, #24, #42, #56,<br>#67, #77, #99, #100,<br>#102 |  |
| *wrky10wrky30*<br>(WK10<sup>+</sup>/WK30<sup>+</sup>) | WK10<sup>ho</sup>/WK30<sup>chi</sup> | 1 | #1 | 15/53.33% |
|  | WK10<sup>bia</sup>/WK30<sup>ho</sup> | 3 | #10, #36, #55 |  |
|  | WK10<sup>bia</sup>/WK30<sup>chi</sup> | 3 | #28, #54, #59 |  |
|  | WK10<sup>he</sup>/WK30<sup>chi</sup> | 3 | #3, #26, #69 |  |
|  | WK10<sup>chi</sup>/WK30<sup>chi</sup> | 5 | #27, #72, #81, #82, #92 |  |

WK10: *VviWRKY10*, WK30: *VviWRKY30*.
Non-transgenic: GFP negative, unedited; Non-edited: GFP positive, unedited.
<sup>+</sup> : Edited, <sup>-</sup> : Unedited, <sup>ho</sup> :Homozygous, <sup>he</sup> :Heterozygous, <sup>bia</sup> :Biallelic, <sup>chi</sup> :Chimeric.
Survival plants are represented by green letters.

### Mutated *VviWRKY10* and/or *VviWRKY30* affect the growth in grapevine

For line 1 in which *VviWRKY10* is homozygously edited (knocked out), compared with WT (the non-edited line 29), it showed narrow leaves and low lignification of stems, and died before transplanting to the matrix (Figure 3A, Table 1). Except for the non-viable plants such as line 1, other gene-edited and control lines were cultured in a growth chamber for two months, then transplanted to the greenhouse and observed for over two years (2021 - 2022). In the growth chamber, the average plant height of *wrky10wrky30* was lower than that of WT, but there was no statistically significant difference between *wrky10* mutants and WT (Figure 3, B and D). After growing in the greenhouse for two years, WT plants grew normally, with a survival rate of 83.72 %; however, among the *wrky10* mutants, all the biallelic mutants died, the heterozygously and chimerically edited plants grew weakly with a total survival rate of 52.83 %; among the *wrky10wrky30* double mutants, three *VviWRKY10* biallelically edited plants survived, of which two were *VviWRKY30* homozygously edited and one was *VviWRKY30* chimerically edited, and the remaining double mutants grew poorly, with a total survival rate of 53.33 % (Table 1). In the greenhouse, all the plant height of mutants was significantly lower than that of WT (Figure 3, C and E). Furthermore, compared with WT plants, ABA content in *wrky10* mutants was significantly decreased, whereas IAA content in *wrky10wrky30* mutants was significantly increased (Supplemental Figure S5).

### *wrky10* single and *wrky10wrky30* double mutants display differential responses to powdery mildew infection

To further study the impact of mutations in *VviWRKY10* and *VviWRKY30* on powdery mildew resistance, some regenerated plants including two *wrky10* lines (line 45 and 70) and two *wrky10wrky30* lines (line 3 and 10) were selected infection tests with *En.* NAFU1, with non-edited WT as control. The inoculated leaves were stained with trypan blue, 3,3’-Diaminobenzidine (DAB), and aniline blue, as shown in Figure 4A, trypan blue staining indicated hypersensitive response (HR)-like cell death induced by *En.* NAFU1 in *wrky10* lines, less in *wrky10wrky30* line 3, while almost no cell death in *VviWRKY30* homozygously edited line 10 and WT; DAB staining and aniline blue staining showed large areas of H_2_O_2_ accumulation and callose deposition in *wrky10* lines, less in *wrky10wrky30* lines, and barely in WT. Compared with WT plants, the average areas showing H_2_O_2_ and callose deposition in *wrky10* increased 55- and 165-fold; conversely, the total hyphal length was reduced to 60% of the WT level. As for *wrky10wrky30* mutant line 10, despite a 20- and 57-fold increase in areas positive for H_2_O_2_ and callose deposition, there was no significant difference in total hyphal length between line 10 and WT (Figure 4, B to D). These results indicate that *wrky10* single mutants displayed strong resistance, whereas the *wrky10wrky30* double mutant line 3 showed only weaker resistance and line 10 with no oblivious resistance.

**Figure 4.**
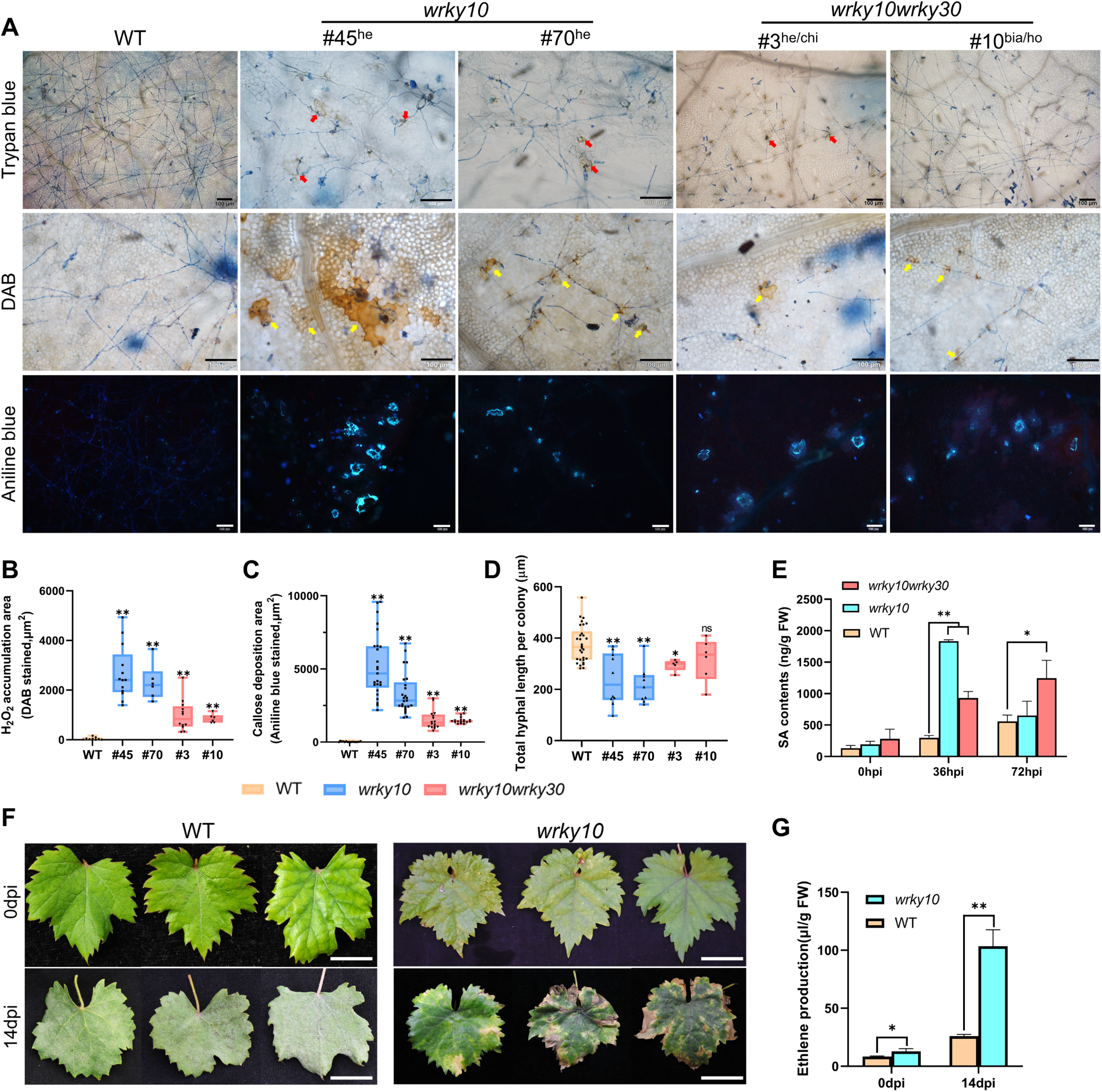
*VviWRKY10* and *VviWRKY30* regulate powdery mildew resistance in grapevine. (**A**) Trypan blue, DAB and Aniline blue staining of WT, *wrky10* and *wrky10wrky30* leaves after inoculation with *En.* NAFU1. The cell death was indicated with red arrows, accumulation of H_2_O_2_ were indicated with yellow arrows. Bars,100 μm. (**B**-**D**) Quantification of H_2_O_2_ accumulation(**B**), callose deposition(**C**) and hyphal length(**D**). Average hyphal length per colony at 3 dpi, accumulation of H_2_O_2_ and callose at 5 dpi. (**E**) SA contents of WT and mutants. Leaves used for detection were collected from WT, *wrky10* and *wrky10wrky30* lines per 36 h post inoculation(hpi). (**F**) Phenotypes of WT and *wrky10* leaves at 0 dpi and 14 dpi. Bars, 2 cm. (**G**) Ethylene production of detached leaves per 36 h which were collected from WT and *wrky10* lines at 0 dpi and 14 dpi. Each data point represents the mean ± standard deviation of three or more biological replicates, and asterisks indicate significant difference compared with WT (Student’s *t* test, **= P<0.01; *= P< 0.05; ns= no significant difference).

To further explore the gene functions of *VviWRKY10* and *VviWRKY30*, *35S:VviWRKY10-eGFP* and *35S:VviWRKY30-eGFP* were constructed and transiently expressed, along with *35S:eGFP* as control, in susceptible grapevine ‘Cabernet Sauvignon’ leaves via *Agrobacterium*-mediated transformation. The transformed leaves were inoculated with powdery mildew and stained with trypan blue at 3 dpi (days post inoculation). As shown in Supplemental Figure S6, the total of hyphal length of OE-WRKY10 was 790 μm, while those of OE-WRKY30 and GFP were 480 μm and 670 μm respectively. These results indicated that overexpression of *VviWRKY30* in leaves increased grapevine resistance to powdery mildew, while overexpression of *VviWRKY10* was just the opposite.

### Mutations in *VviWRKY10 and VviWRKY30* affect gene expression in multiple defense pathways

The activation of SA, ET and ROS pathways induced by pathogens plays a key role in plant defense. To explore changes in these pathways in mutants, some key genes were selected and measured by qRT-PCR assays in the mutant and WT leaves. As shown in Figure 5, in *wrky10*, *VviPAD4* was highly induced by *En.* NAFU1 at 48 hpi, which is 5-fold of the WT expression level, while *VviEDS1, VviEDS5-2*, *VviPR1*, *VviNPR2, VviACS3*, *VviACS3L* and *VviERF3* were always more highly expressed upon inoculation, with their peak values being 40-, 14-, 200-, 6-, 197-, 32- and 4.7-fold of the expression levels of the WT respectively. The transcription level of *VviACS2* in *wrky10* was 2.5-fold of that in WT at 0 hpi, and there was no significant difference after inoculation, even lower than WT at 36 hpi, while expression of *VviEIN2* and *VviRBOHD2* peaked at 48 and 36 hpi, showing 2.6- and 9.3-fold increase compared to that in WT, respectively. In *wrky10wrky30*, the transcription levels of *VviPAD4*, *VviNPR2*, *VviACS2* fluctuated at different time points but over all there was no significant difference compared to those of WT. Interestingly, *VviEDS1, VviEDS5-2*, *VviPR1*, *VviICS2*, *VviERF3* and *VviRBOHD2* in *wrky10wrky30* exhibited higher levels upon inoculation, with their peak values being 19-, 12-, 270-, 9-, 4.8- and 7.2-fold of the levels in WT respectively. The expression of *VviACS3* in *wrky10wrky30* was 51-fold of that in WT at 0 hpi, but soon returned to the WT level after inoculation, while there was no significant difference in the expression of *VviACS3L* and *VviEIN2* between *wrky10wrky30* and WT. Collectively, the above results showed that most of the genes involved in the SA, ET and ROS pathways in *wrky10wrky30* and *wrky10* in particular were significantly increased upon inoculation with *En.* NAFU1, which may explain the enhanced disease resistance in these mutant lines especially *wrky10*.

**Figure 5.**
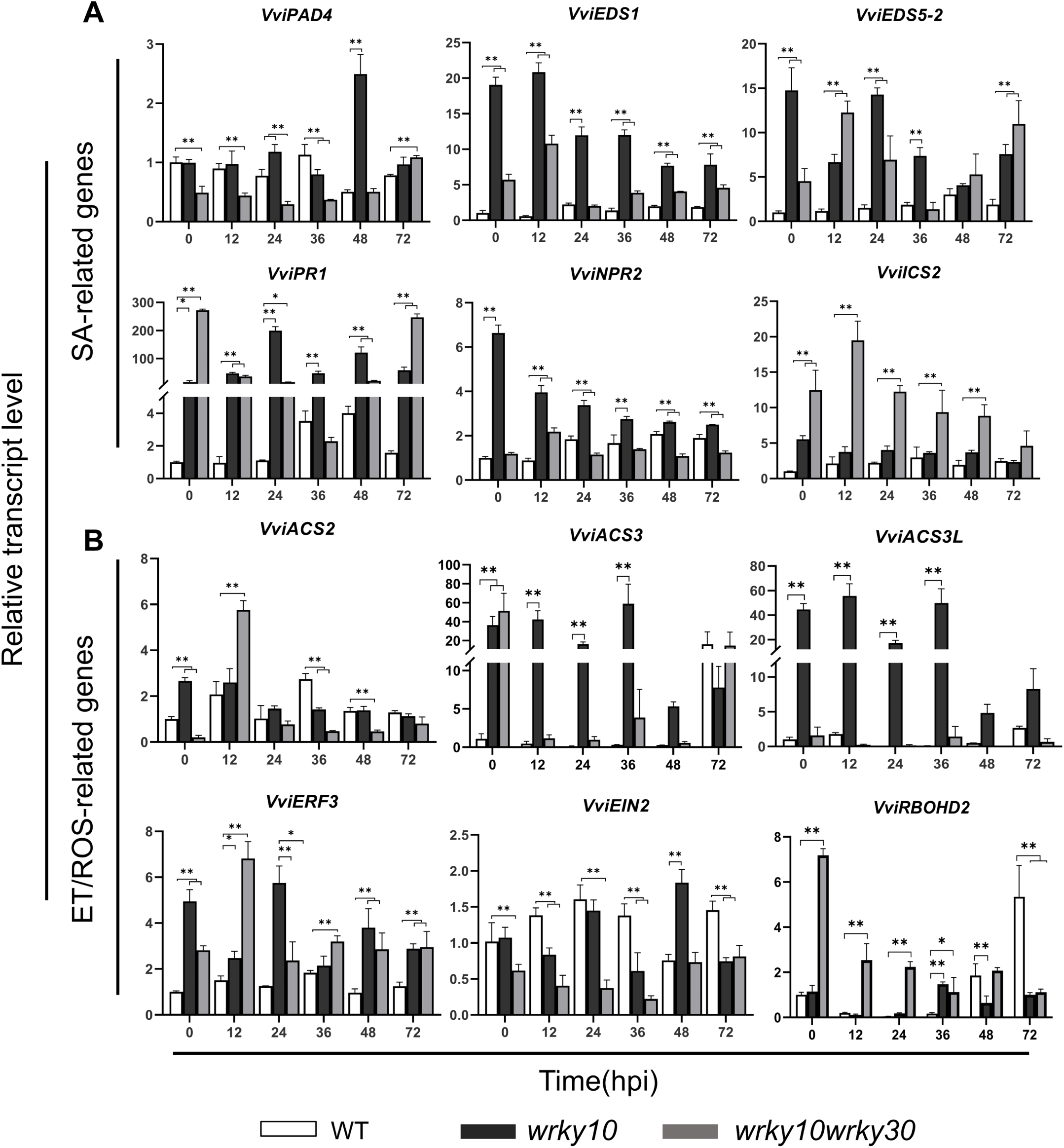
The expression of defense-related genes in WT and mutant lines post inoculation of *En.* NAFU1. (**A**) Relative transcript level of SA-related genes. (**B**) Relative transcript level of ET/ROS-related genes. *VviACTIN7* gene was used as an endogenous control. Each data point represents the mean ± standard deviation of three biological replicates, and asterisks indicate significant difference compared with WT(GenBank accession numbers: *VviPAD4* XM_010654614.2, *VviEDS1* NM_001281038.1, VviEDS5-2 XM_002274777.4, *VviPR1* XM002273752, *VviNPR2* XM_002274009.3*, VviICS2* XM_019226638.1*, VviACS2* XM_002278453.4*, VviACS3* XM_003635528.3*, VviACS3L* XM_002269744.4*, VviERF3* XM_002277986.3*, VviEIN2* XM_002276363.3*, VviRBOHD2* XM_019222717.1.) (Two-way ANOVA, **= P<0.01; *= P< 0.05).

To further examine if the upregulation of the pathway genes indeed leads to increases biosynthesis of SA or ET in the mutant line, content of SA and ET in leaf tissues of WT and the mutant lines was measured prior to, 36 hpi, 72 hpi and 14 dpi (for ET only). As expected, SA content of *wrky10* and *wrky10wrky30* were 6- and 3-fold of that of WT respectively at 36 hpi; SA content of *wrky10* dropped to the level of the WT and maintained at a low level, while SA content of *wrky10wrky30* was still higher than that of the WT (Figure 4E). Because leaves of *wrky10* infected with powdery mildew (but not those of *wrky10wrky30*), often senesced earlier than those of WT (Figure 4F, Supplemental Figure S7), ET content in *wrky10* and WT (but not the *wrky10wrky30* double mutant because of their weak growth and insufficient leaf tissue for analysis; Figure 3C) was measured prior to and at 14 dpi with *En.* NAFU1. The ET content in *wrky10* was slightly higher (1.5-fold) than that in WT in the absence of *En.* NAFU1; however, the ET content in *wrky10* further increased significantly upon infection by powdery mildew, reaching ∼8x of that in WT at 14 dpi (Figure 4G). These results suggested that VviWRKY10 and VviWRKY30 affected the resistance to powdery mildew by changing the accumulation of SA and ET in grapevine.

### VviWRKY10 and VviWRKY30 regulate the transcription of SA-, ET- and ROS-related genes

WRKY TFs function mainly by binding to the W-box (TTGACC/T), a typical cis-acting element on the promoters of the target genes (Rushton et al., 2010). We selected genes of the SA, ET and ROS pathways, analyzed their promoters, and found that all of them contained at least one canonical W-box (Supplemental Figure S8A). Further, DLUC reporter assays was used to identify the relationship between VviWRKY10, VviWRKY30 and potential downstream target genes. Different combinations of reporters and effecters, in which *35S:GFP* was used as the control of the effecter and *35S:REN* (*Renilla luciferase*) was used as an internal control (Figure 6A), were used to co-transform tobacco protoplasts, followed by measuring LUC (Firefly luciferase) and REN activities sequentially to reflect the transcriptional activity of individual promoters *in vivo*. As shown in Figure 6B, compared to GFP, VviWRKY10 and VviWRKY30 inhibited promoter activities of six SA-related genes (*VviICS2, VviEDS5-1, VviEDS5-2, VviPR1, VviPR5* and *VviPR10.3L2*), two ET-related genes (*VviACS1* and *VviEIN2*), and one ROS-related gene (*VviRBOHD2*), while they had no significant effect on the promoter activity of *VviACS2*. Additionally, VviWRKY30 significantly enhanced promoter activities of *VviNPR2*, *VviACS3* and *VviACS3L*. The results suggest that VviWRKY10 and VviWRKY30 could inhibit the promoter activities of multiple genes involved in the SA, ET and/or ROS pathways and that VviWRKY30 may also enhance the expression of other genes in the same pathways.

**Figure 6.**
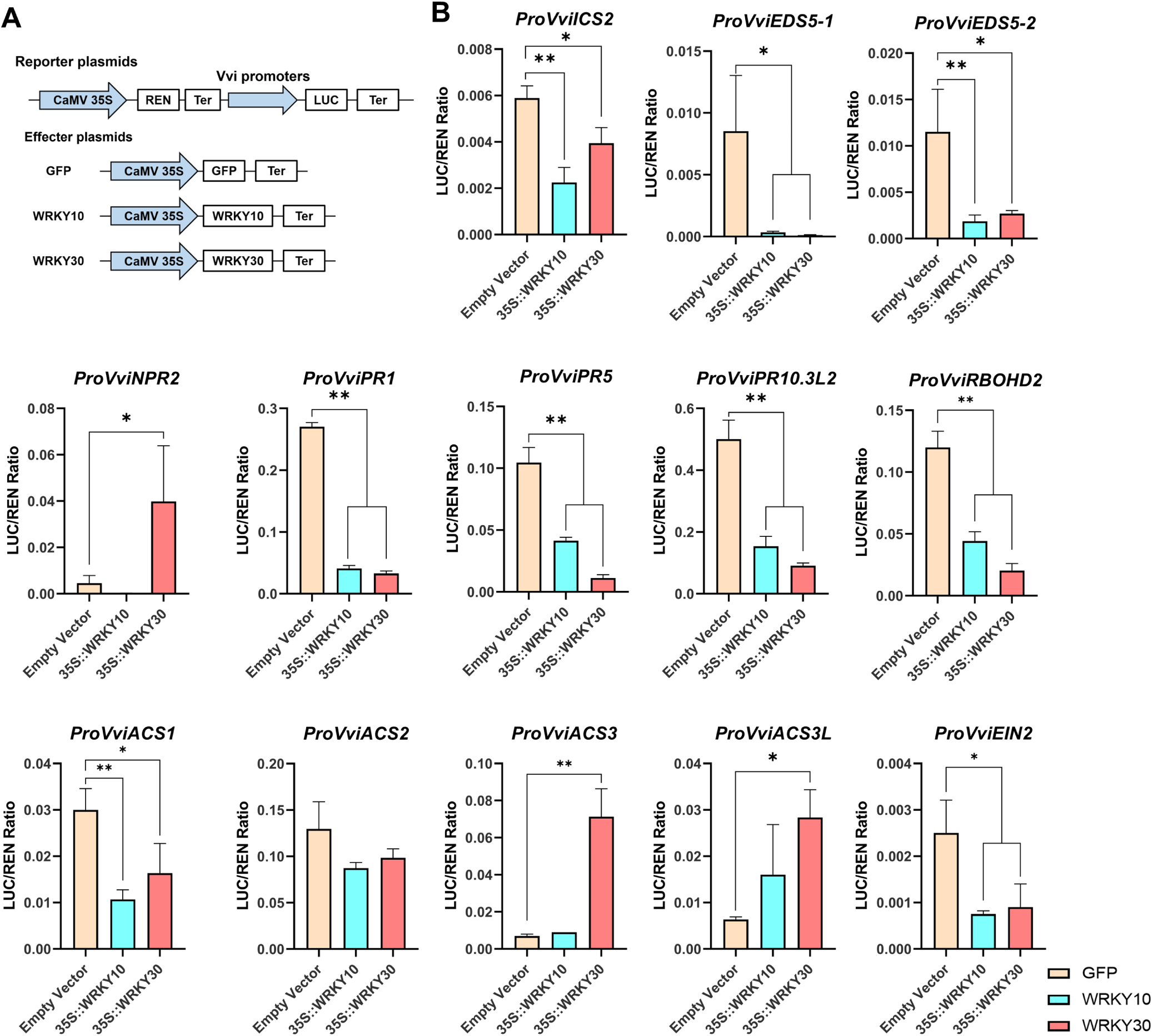
VviWRKY10 and VviWRKY30 regulate the transcription of SA, ET and ROS related genes by dual LUC reporter assays. (**A**) Schematic diagram of reporter and effecter constructs used in the DLUC reporter assays. In the reporter plasmids, Renilla (REN) luciferase gene driven by CaMV35S promoter was used as the internal control, and the target gene promoters are fused to the *LUC* (*Firefly Luciferase*). In the effecter plasmids, *GFP* (control), *WRKY10* and *WRKY30* are driven by CaMV35S promoter. Ter, transcriptional terminator sequence. (**B**) Dual luciferase reporter assays results. Different combinations of reporter and effecter plasmids were co-expressed in protoplast of *N.benthamiana*. The ability of WRKY10 and WRKY30 to regulate the reporter LUC gene was represented by the ratios of LUC to REN. The *35S:REN* served as an internal control. Each data point represents the mean ± standard deviation of three biological replicates, and asterisks indicate significant difference compared with GFP(Student’s *t* test, **= P<0.01; *= P< 0.05).

### VviWRKY10 and VviWRKY30 directly bind to the promoters of SA-, ET- and ROS-related target genes

To verify whether *VviWRKY10* and *VviWRKY30* directly regulate SA, ET and ROS pathways via binding to the W-boxes in the promoters of the genes whose transcription was inhibited or activated as shown above, we selected six genes *VviEDS5-2, VviPR1, VviPR5, VviACS3, VviACS3L* and *VviRBOHD2* for the tests by using ChIP-qPCR and EMSA. The probes and primers used in the experiment were labeled on each gene promoter (Supplemental Figure S8B). After analyzing the cloned genes, it was found that the predicted protein sequences of VviACS3 and VviACS3L were highly conserved, differing only in one amino acid (Supplemental Figure S9) in addition to their exact same W-box sequences in their promoters. Hence, the same probe and primers were used to detect *VviACS3* and *VviACS3L* (Supplemental Figure S8B, Supplemental Figure S9A).

First, we used grapevine callus overexpressing *VviWRKY10* or *VviWRKY30* for ChIP assays. As shown in Figure 7, A to E, VviWRKY10 and VviWRKY30 were significantly enriched in fragments including F2 of *ProEDS5-2*, F1 of *ProPR1*, F2 of *ProPR5*, F2 of *ProACS3*/*ACS3L*, F1 and F4 of *ProRBOHD2*. In addition, VviWRKY10 was enriched to F3 of *ProRBOHD2*; VviWRKY30 was enriched to F1 of *ProEDS5-2*, F2 of *ProPR1*, and F2 of *ProRBOHD2*. Next, His-tagged VviWRKY10 and VviWRKY30 were expressed in *E. coli* and the purified proteins together with W-box-containing oligonucleotide probes were subjected to EMSA, with unlabeled and mutant probes as competitors. The results indicated that the purified VviWRKY10 or VviWRKY30 proteins could specifically bind to the W-box (Supplemental Figure S10). Then, similar EMSA was performed using purified VviWRKY10 or VviWRKY30 proteins and labeled oligonucleotide probes synthesized based on promoter sequences enriched by both of these two proteins, with unlabeled sequences as competitors. The results showed that both VviWRKY10 and VviWRKY30 could form protein-probe complex hysteresis bands with the labeled probes, and the bands were significantly weakened when the unlabeled probes were present (Figure 7, F to J). Thus, results from our *in vivo* and *in vitro* experiments consistently demonstrated that VviWRKY10 and VviWRKY30 could directly bind to the promoters of *VviEDS5-2*, *VviPR1*, *VviPR5*, *VviACS3*/*ACS3L*, and *VviRBOHD2*.

**Figure 7.**
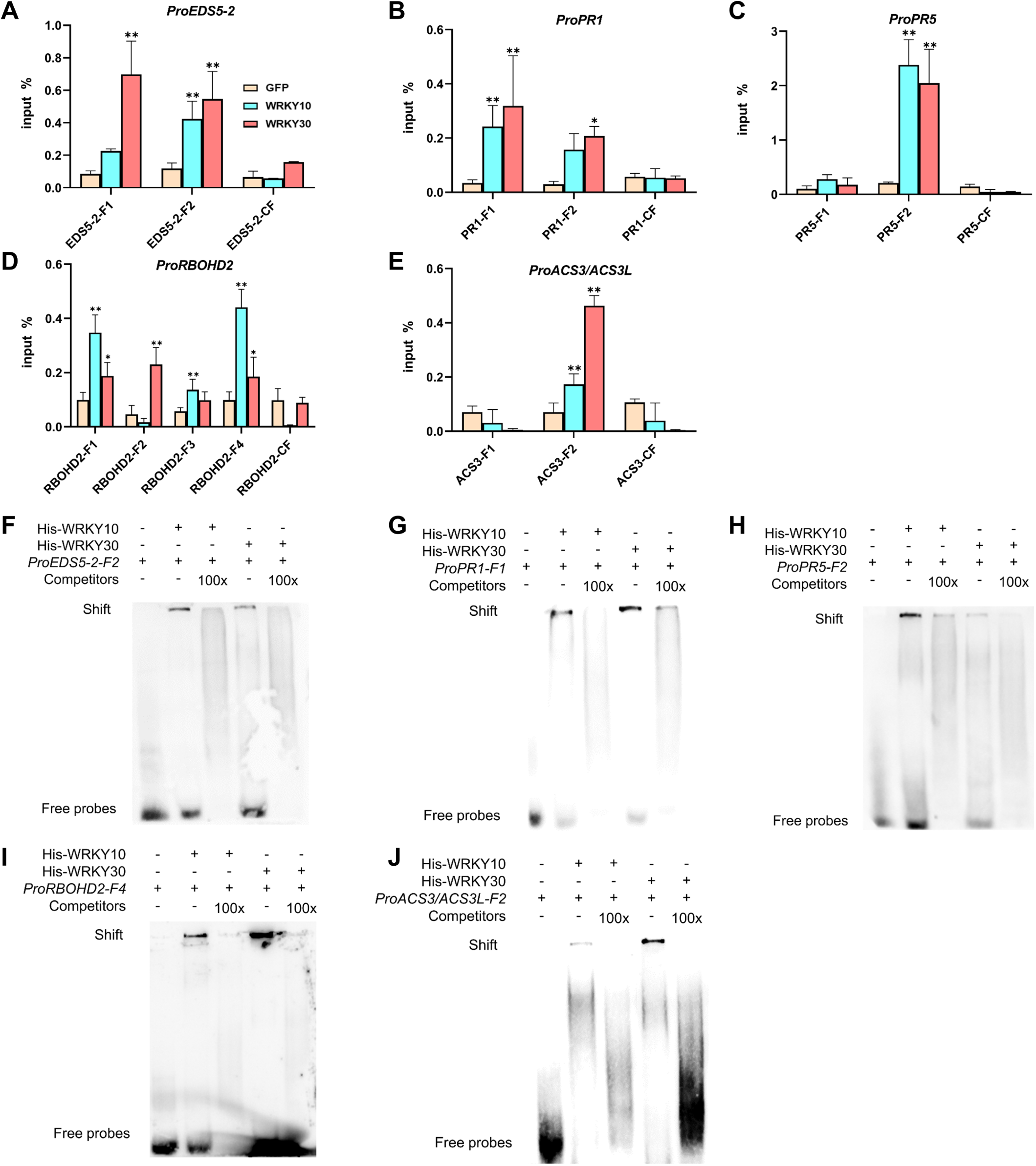
Chromatin immunoprecipitation (ChIP)-qPCR and Electrophoretic mobility shift assays (EMSA) to examine the association between WRKY10, WRKY30 and its targets. (**A-E**) Association of WRKY10 and WRKY30 with its targets by ChIP-qPCR assays. Chromatin prepared from WRKYs-GFP callus were detected with qPCR with chromatins prepared from GFP as the control. Each data point represents the mean ± standard deviation of three biological replicates, and asterisks indicate significant difference compared with CF (F1-F4, sequences of fragments designed from 5’ to 3’ WRKYs may bind; CF, fragments that WRKYs do not bind). (**F-J**) Competitive EMSA to detect the binding of WRKY10 and WRKY30 to promoters of target genes.*VviACS3* and *VviACS3L* were used the same fragments (Supplemental Figure S8). The DNA binding assays were performed using purified His-WRKYs protein and biotin-labelled fragments of the promoters containing the W-boxes, using non-labeled fragments as competitors in a molar excess of 100x. The + and - symbols indicate the presence and absence of components. The bands at the upper and lower part of membranes indicate shift (protein-probe complex) and unbound free probes, respectively (Student’s *t* test, **= P<0.01; *= P< 0.05).

### VviWRKY10 and VviWRKY30 directly bind to mutual promoters and inhibit expression

It was reported that AtWRKY18 and AtWRKY40 physically associate with each other in *Arabidopsis thaliana* (Xu et al., 2006). In order to explore whether VviWRKY10 and VviWRKY30 interact with each, we first measured the mRNA levels of *VviWRKY30* in leaves of *wrky10* and WT prior to and post-inoculation with *En.* NAFU1. As shown in Figure 8A, at 0 hpi, the level of *VviWRKY30* in *wrky10* was 51-fold than that of WT. Although the transcript level decreased after inoculation, it was still higher than that of WT at most time points (Figure 8A). Further, LUC assays showed that VviWRKY10 can inhibit its own and *VviWRKY30* promoter activities, which can be partially verified by the increased expression of *VviWRKY30* in *wrky10* (Figure 8, A and B). In addition, VviWRKY30 can also inhibit the promoter activity of *VviWRKY10* (Figure 8C). Finally, EMSA and ChIP assays further showed that VviWRKY10 and VviWRKY30 could directly bind to each other’s promoters (Figure 8, D to G). These results suggested that VviWRKY10 and VviWRKY30 may be involved in mutual inhibition of each other’s transcription.

**Figure 8.**
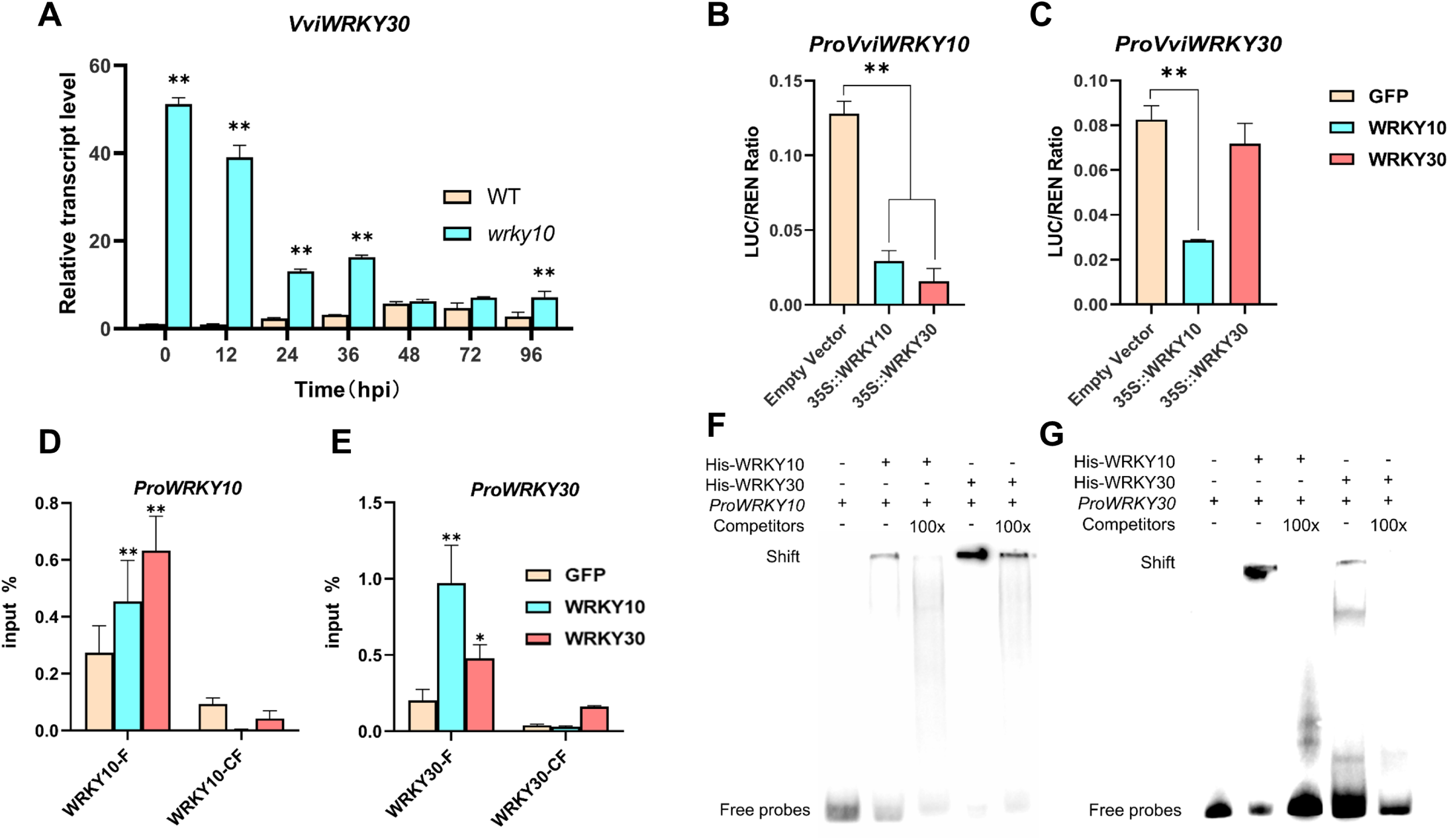
VviWRKY10 and VviWRKY30 directly bind to mutual promoters and inhibit expression. (**A**) Relative transcript level of *VviWRKY30* in WT and *wrky10*. *VviACTIN7* gene was used as an endogenous control. (**B**) WRKY10 and WRKY30 repress the expression of *VviWRKY10* by dual LUC reporter assays. (**C**) WRKY10 repress the expression of *VviWRKY30* by DLUC. (**D**-**G**) EMSA and ChIP assays of WRKY10 and WRKY30 binding to each other promoters. Each data point represents the mean ± standard deviation of three biological replicates, and asterisks indicate significant difference compared with WT(**A**), GFP(**B,C**), and CF(**D,E**) (F, sequences of fragments designed from 5’ to 3’ WRKYs may bind; CF, fragments that WRKYs do not bind; two-way ANOVA(**A**), student’s *t* test(**B-E**), **= P<0.01; *= P< 0.05).

## DISCUSSION

WRKY transcription factors play important regulatory roles in defense against pathogens in plants. The Arabidopsis WRKY transcription factors, *AtWRKY18* and *AtWRKY40*, act redundantly as negative regulators of basal resistance against powdery mildew in Arabidopsis (Chen & Chen, 2002, Xu et al., 2006, Shen et al., 2007, Pandey et al., 2010, Abeysinghe et al., 2019). In this study, we investigated the roles of two grapevine transcription factors, VviWRKY10 and VviWRKY30, as probably orthologs of AtWRKY18 and AtWRKY40, in grapevine defense responses against powdery mildew. Our results have revealed complex regulatory mechanisms of these two grapevine transcription factors that are distinct from the relatively simple functional redundancy of their Arabidopsis orthologs.

In an early study, Pandey et al. proposed that the adapted powdery mildew pathogen *Golovinomyces orontii* alters the balance of the SA-pathway and the JA-pathway via impacting the functionally redundant AtWRKY18/40 in Arabidopsis during early infection, thereby subverting host defense (Pandey et al., 2010). Our study found that *VviWRKY10* and *VviWRKY30* may play distinct roles in different stages of *E. necator* infection. *VviWRKY10* was induced in an early stage of infection, while *VviWRKY30* was induced at a later time, implying differential roles in transcriptional regulation of host defense (Figure 1, B and C). Consistently, *E. necator*-induced ROS and callose accumulation in *wrky10* were significantly increased compared with those in the WT, which correlates with a significant reduction of hyphal growth in leaves of *wrky10* compared to that of the WT (Figure 4, A to D), supporting a role of *VviWRKY10* in negative regulation of defense against powdery mildew. Interestingly, even though in the *wrky10wrky30* double mutant also showed enhanced resistance to the pathogen, the level of resistance was lower than that of *wrky10*, and there was considerable mycelium growth (Figure 4, Table 1), implying a complex and perhaps antagonistic relationship between VviWRKY10 and VviWRKY30. Indeed, while transient overexpression of VviWRKY10 in grapevine leaves led to significantly increased hyphal length of *En*. NAFU1, transient overexpression of VviWRKY30 did the opposite (Supplemental Figure S6), further suggesting that VviWRKY10 and VviWRKY30 play opposing roles in modulating defense against powdery mildew in grapevine.

Given that plants adopt contrasting strategies to fight against biotrophic and necrotrophic pathogens, our results on *VviWRKY10* are consistent with and provide further explanations for the earlier observation that overexpression of *VaWRKY10* in *V. vinifera* cv. Thompson Seedless significantly improved resistance to *B. cinerea*, a necrotrophic fungal pathogen (Wan et al., 2021). Specifically, we found that the enhanced resistance to powdery mildew in the *wrky10* single mutant was associated with the activation of SA, ET and ROS pathways (Figure 4, Figure 5).

Notably, our results on *VviWRKY30* also largely agree with the results of an earlier study where *VviWRKY30* (named as *VviWRKY40* in that study) was reported to play a role in PvRXLR111-mediated suppression of flg22-induced ROS production, thereby promoting *Phytophthora capsici* infection (Ma et al., 2021). Our study provides more mechanistic insight onto how VviWRKY30 works: VviWRKY30 binds to the *VviRBOHD2* promoter and inhibit its expression. This conclusion was supported by our genetic evidence that loss of VviWRKY30 in the *wrky10wrky30* double mutant resulted in increased *VviRBOHD2* expression (Figure 5B, Figure 6B, Figure 7, D and I). Furthermore, we found that increased expression of *VviWRKY30* in the *wrky10* mutant may be attributable to the increased expression of *VviACS3* and *VviACS3L* genes, thereby activating the ET pathway (Figure 6B, Figure 7, E and J), which in turn may partially explains the enhanced resistance to *En*. NAFU1 at later stages of infection (Figure 4G, Figure 5B, Figure 8A).

It is well known that WRKYs usually play roles in transcriptional regulation of plant defense (Eulgem & Somssich, 2007). For example, it has been shown that AtWRKY18 and AtWRKY40 directly bind to the promoters of multiple genes acting in the SA, ET and JA pathways, directly regulating their transcription, resulting in the formation of a complex regulatory network (Birkenbihl et al., 2017). In this study, using DLUC, ChIP and EMSA assays, we demonstrated that VviWRKY10 can directly bind to the promoters of *EDS5-2*, *PR1*, *PR5*, *RBOHD2* and *WRKY30* and inhibit the expression of these target genes, thereby inhibiting the accumulation of SA and ROS, and the WRKY30 protein (Figure 6B, Figure 7, A to D, F to I, Figure 8, C, E and G). Intriguingly, we also showed that VviWRKY30 can directly bind to the promoter of *VviWRKY10*, inhibiting its expression and downregulating the SA pathway at an early stage of infection while promoting the ET pathway for limiting infection at a later stage (Figure 6B, Figure 7, E and J, Figure 8, B, D and F).

Based on the results from this study, we propose a working model for the role of VviWRKY10 and VviWRKY30 in the regulation of powdery mildew resistance in grapevine (Figure 9). Specifically, we propose: (i) VviWRKY10 inhibits the expression of *VviEDS5-2, VviPR1, VviPR5,* and *VviRBOHD2*, presumably to avoid overactivation of the SA-dependent defense responses upon powdery mildew infection in grapevine; (ii) VviWRKY30 promotes the expression of *VviACS3* and *VviACS3L* to increase ET production for limiting powdery mildew growth at a later stage of infection; and (iii) VviWRKY10 can inhibit VviWRKY30 accumulation to inhibit ET synthesis, while VviWRKY30 can also inhibit VviWRKY10 accumulation to promote production of SA and ROS. It is possible that the distinct roles of VviWRKY10 and VviWRKY30 together with their mutual inhibition are required for the activation of measured and balanced defenses involving multiple hormonal pathways against powdery mildew and likely other pathogens in grapevine.

**Figure 9.**
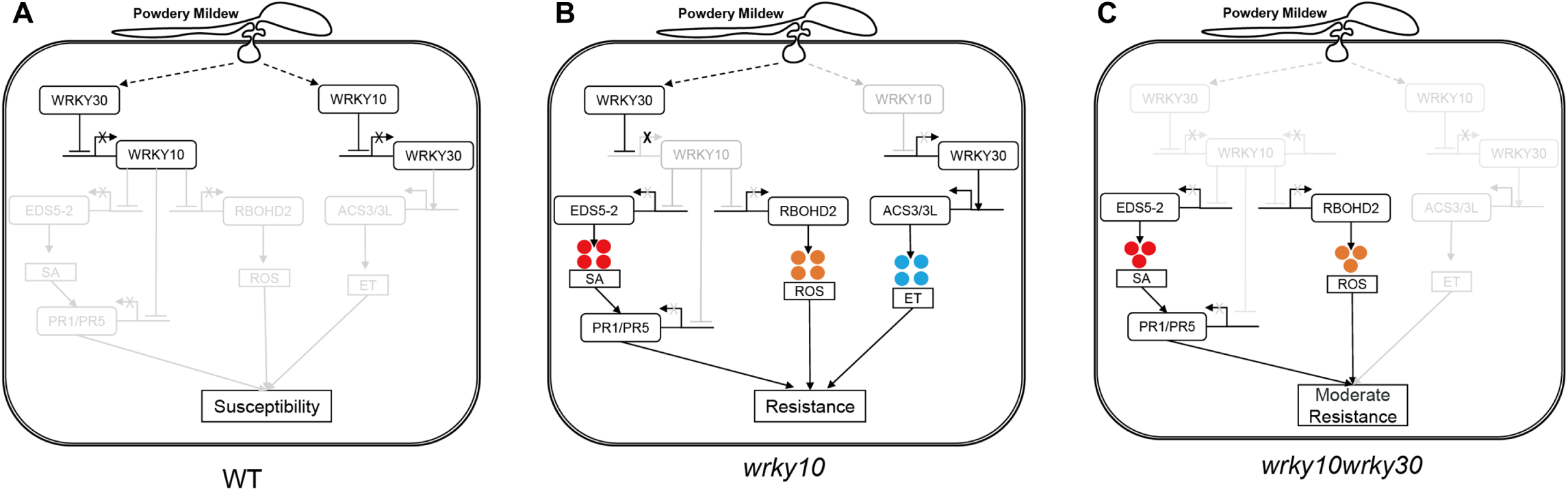
A proposed work model of two grapevine transcription factors (WRKY10 and WRKY30) in regulating powdery mildew resistance. (**A**) In WT, WRKY10 and WRKY30 inhibited each other and both downstream pathways were inhibited, susceptible to powdery mildew. (**B**) In *wrky10*, the SA and ROS pathways were activated, and WRKY30 activated the ET pathway, resulting in resistance to powdery mildew. (**C**) In *wrky10wrky30*, only SA and ROS pathways were activated, resulting in moderate resistance to powdery mildew.

## Materials and methods

### Plant materials and growth conditions

The WT *Vitis vinifera* cv. Cabernet Sauvignon and transgenic plantlets were cultured under the same conditions. In April and May of 2021, we observed the phenotypes of WT, *wrky10*, and *wrky10wrky30* mutant plants in chambers with temperature ranging from 22 to 27 °C, relative humidity ranging from 75 % to 90 %, and under a long-day photoperiod (16 h : 8 h, light (200 μmol m^-2^ s^-1^) : dark). From May 2021 to November 2023, the WT and mutant plants were transplanted in a greenhouse with temperature ranging from 25 to 35 °C (in summer) or 5 to 25 °C (in winter), relative humidity ranging from 40 % to 75 %.

### Cloning and sequence analysis

The *VviWRKY10* and *VviWRKY30* genes were amplified from ‘Cabernet Sauvignon’ leaf cDNA using Planta Max Super-Fidelity DNA Polymerase (Vazyme Bio Co., Nanjing, China). The amplified sequences were integrated into the pMD-19T vector and then introduced into *E. coli* DH5α. Ten clones of each gene were randomly selected and sequenced. The amplification primers were designed according to the genome database of *V. vinifera* cv. Pinot Noir in the NCBI database (https://www.ncbi.nlm.nih.gov/). The sequences of amplified primers are shown in Supplemental Table S1.

The coding sequences of *VviWRKY10* and *VviWRKY30* genes obtained by sequencing were translated into protein sequences by DANMAN, and then BLAST-P search was performed in NCBI. The genes with the highest similarity to *VviWRKY10* and *VviWRKY30* in other species were selected, and the phylogenetic tree was constructed by MEGA-X. The parameters were set as follows: bootstrap value was 1000, p-distance model was selected, partial deletion value was 50, and other default settings. The protein sequences of VviWRKY10, VviWRKY30 and AtWRKY18, AtWRKY40 and AtWRKY60 were analyzed by Jalview, and the parameters were set by default.

### Subcellular location assays

The coding sequences of *VviWRKY10* and *VviWRKY30* were amplified and ligated into the pCAMBIA2300 vector, which contains a 35S promoter and a C-terminal GFP tag. The recombinant vectors and *35S:AtH2B-mCherry* fusion expression vector were transformed into tobacco leaves respectively combined. Fluorescence photos were taken using a confocal laser scanning microscope (LEICA TCS SP8, Germany) 3 days after transfection.

### Target selection and vector construction

In the genome of ‘Pinot Noir’, *VviWRKY10* gene is located on chromosome 4, 2201 bp in length, encoding 296 amino acids. *VviWRKY30* gene is located on chromosome 9, the full length of 1561 bp, encoding 311 amino acids. Select target points according to design principles (Xing et al., 2014). Based on the NCBI database, BLAST-P search was performed on the target sequences using the ‘Pinot Noir’ genome to ensure the specificity of the target sites selection. At the same time, the amplified *VviWRKY10* and *VviWRKY30* were compared to ensure that the selected target sites were completely consistent in ‘Cabernet Sauvignon’. According to the sequence of vector pKSE401-GFP and intermediate vector pCBC-DT1T2, specific primers containing *Bsa* I restriction site were designed. The pCBC-DT1T2 plasmid was used as a template for PCR amplification using Planta Max Super-Fidelity DNA Polymerase to obtain two sgRNA expression cassettes. The purified PCR products were assembled into the pKSE401-GFP vector. The primers are shown in Supplemental Table S1.

### Plant transformation and detection of mutations

The grapevine PEM was transformed by *Agrobacterium*-mediated genetic transformation (Wan et al., 2020). Using stereomicroscope (MZ10F, LEICA, Germany) to observe whether the plantlets with GFP fluorescence, screening transgenic positive plants. The 0.5-1.0 g leaves of regenerated positive ’Cabernet Sauvignon’ plantlets were taken, and the whole gDNA was extracted by CTAB method. Gene-specific primers were used to amplified the positive plants across the target region and recovered the PCR products for sequencing. DNA sequence alignment was performed on the sequencing results to determine whether editing occurred, and then the editing type was analyzed by ’DSDecodeM’ (Liu et al., 2015). For the edited plants whose ’DSDecodeM’ decoding failed, the PCR purified product was cloned into pMD19-T, and 10 single clones were randomly selected for Sanger sequencing to analyze the editing type. Multiple sequence alignment and amino acid sequence analysis were performed using Jalview to determine the effect of each gene mutation type on amino acids. The primers are shown in Supplemental Table S1.

### Evaluation of resistance to powdery mildew

In order to explore the phenotype of *wrky10* single mutant and *wrky10wrky30* double mutant after powdery mildew infection, the leaves of the mutants and WT were inoculated with *En*. NAFU1 (Gao et al., 2016). The leaves were stained with trypan blue at 3 dpi, DAB and aniline blue at 5 dpi to observe the growth of powdery mildew and the cell reaction of grapevine leaves (Hu et al., 2018, 2019). At the same time, the inoculated leaves were frozen for qRT-PCR and hormone measuring after inoculation.

### *Agrobacterium*-mediated transient expression in grapevine leaves

For transient overexpression of *VviWRKY10* and *VviWRKY30*, the coding sequences of these two genes were translationally in-frame fused with eGFP in the binary vector pCAMBIA2300 under control of the *35S* promoter via homologous recombination. The resulting *35S:VviWRKY10-eGFP* and *35S:VviWRKY30-eGFP*, as well as *35S:eGFP* DNA constructs were introduced into *Agrobacterium* tumefaciens strain GV3101. Agrobacterial cells were grown to OD_600_=0.5-0.6, spun down at 4000 rpm for 5 min, and resuspended in an equal volume of buffer (0.5 % (w/v) glucose, 50 mM MES (pH 5.6), 3 mM Na_2_HPO_4_, 100 μM acetosyringone). After incubation at 28 ℃ for 1 h, the resuspended bacterial solution was injected via the leaf abaxial side into the third to fifth fully expanded ‘Cabernet Sauvignon’ leaves. Two days later, the infiltrated leaves were inoculated with *En.* NAFU1. At 3 dpi, Trypan blue was used to visualize fungal structures and hyphal lengths were measured with the aid of an Olympus BX-63 microscope (Japan).

### RNA extraction and quantitative real-time PCR (qRT-PCR) assays

The leaves of ’Cabernet Sauvignon’ were inoculated with *En*. NAFU1. The leaves were collected at 0 h, 12 h, 24 h, 36 h, 48 h, 72 h, and 96 h post inoculation. At 12 hpi, parts of 5-7 inoculated leaves were randomly cut for RNA sample collection, and trypan blue staining was performed to check spore germination. After spore germination, 5 leaf pieces (∼100 mg) were randomly collected at each time point as one leaf sample for RNA extraction using the E.Z.N.A. Plant RNA Kit (Omega, Guangzhou, China). About 1 μg RNA was used for cDNA synthesis using HiScript Q Select RT SuperMix (Vazyme, Nanjing, China). The cDNA samples served as template of qPCR for measuring mRNA levels of the selected defense genes and *VviACTIN7* (XM_002282480.4) as internal control. The relative transcription level of the defense genes was calculated with the 2^-△△ Ct^ method. The expression level for each gene was presented as the mean value of three biological replicates, and significance testing was performed by two-way analysis of variance. The sequences of the primers used for qRT-PCR were listed in Supplemental Table S1.

### Hormone measurements

Taken leaves at 0, 36, and 48 hpi, 0.1 g was weighed and put into 2 mL sterile centrifuge tubes, sterilized steel beads were added, and quickly frozen in liquid nitrogen. The leaves were fully ground with a tissue grinding instrument, and 1 mL of ethyl acetate extract was added. After fully mixing for 10 min, centrifuged at 10000 rpm, 4 ℃ for 5 min, the supernatant was transferred to new centrifuge tubes. The organic phase in the centrifuge tube was blown dry with a nitrogen blowing instrument, and 200 μL 50 % methanol solution was added to fully shake and mix. After centrifugation at 10000 rpm for 5 min, the supernatant was absorbed with a 1 mL syringe, and filtered into a liquid injection bottle using 0.22 μm organic filters.

The leaves at 0 dpi and 14 dpi were made into leaf discs. 20 leaves in each group were randomly selected in 10 mL headspace bottles, sealed in the incubator for 36 h, and 1 mL of gas was extracted to determine ethylene by gas chromatography. was calculated by standard curve method.

The quantitative analysis of each hormone and ethylene content were performed using the standard curve method.

### Promoter analysis and dual luciferase reporter assays (DLUC)

The DNA sequence of 2000 bp upstream of the start codon of *VviWRKY10*, *VviWRKY30*, SA-, ET-, and ROS-related genes were selected for DLUC assays. Gene Regulation (gene-regulation.com) and PlantCARE, a database of plant promoters and their *cis*-acting regulatory elements (bioinformatics.psb.ugent.be/webtools/plantcare) were used to predict *cis*-acting elements such as W-boxes from the selected promoter sequences. The diagrams showing the *cis*-acting elements of the analyzed genes were drawn by Gene Structure Display Server (gsds.gao-lab.org).

The coding sequences of *VviWRKY10* and *VviWRKY30* were amplified and ligated into the pBI221-GFP vector as effecters, the promoters of defense-related gene were inserted into the 35S-Rluc-35S-Fluc vector as reporters. According to the previous method, different combinations of reporter and effecter plasmids were co-expressed in tobacco protoplast (Zhao et al., 2016). Full-wavelength microplate reader (Infinite M200pro, Tecan, Switzerland) for detection, and calculation of relative transcriptional activity based on LUC to REN ratio.

### Electrophoretic mobility shift assays (EMSA)

The coding sequences of *VviWRKY10* and *VviWRKY30* were amplified and cloned into pET30a vector. The His-WRKY10 and His-WRKY30 fusion protein were expressed in *E.coli* strain Rosetta (DE3). The fusion protein was induced by 0.2 mM IPTG and purified by protein purification instrument (NGC Discover 10). Complete the EMSA experiment according to the Chemiluminescent EMSA Kit instructions (Beyotime, China). Labeled probes, unlabeled probes and mutant probes sequences are shown in Supplemental Table S1.

### Chromatin immunoprecipitation (ChIP)-qPCR assays

The grapevine transgenic callus stably expressing VviWRKY10-GFP or VviWRKY30-GFP were used as materials. 1 g callus was cross-linked with 1% formaldehyde for 15 min, and then glycine with a final concentration of 100 mM was added to terminate the reaction, washed twice with distilled water and frozen in liquid nitrogen. Next, the chromatin was treated with ultrasound and the DNA fragments were precipitated using GFP-TRAP Magnetic Agarose beads (Chromotek, gtma-20). After protein digestion, the precipitated DNA was purified and directly used as qPCR template. The transformed GFP empty callus was used as the negative control, and the DNA without magnetic beads precipitation was used as input. The results are presented as a percentage of input. Sequences at least 600 bp after the translation initiation site were used as controls. The primers are shown in Supplemental Table S1.

### Statistical analysis

All variance tests were performed using GraphPad Prism 8. In this paper, two-way ANOVA was used for all gene expression analysis (Figure 1B, C, Figure 5, Figure 8A) and Student’s *t* test was used for the rest experiments. Asterisks above columns indicate significant differences (**= P < 0.01; *= P < 0.05; ns= no significant difference).

## Acknowledgments

We thank Dr. Wei Rong of Hainan University for kindly providing pET30a vector, and Dr. Kunming Chen of Northwest A&F University for kindly providing 35S-Rluc-35S-Fluc vector. This work was supported by the National Natural Science Foundation of China (Grant No. 32272670, 31972986) to Y.Q.W. and National Science Foundation (Grant No. IOS-1901566 to S. X.) The authors would like to thank the anonymous reviewers for their comments on the manuscript.

## Author contributions

Y.Q.W. conceived the study with help from S.X.; M.Z., H.Y.W. performed the experiments; X.N.Y., K.C.C. and Y.H. conducted data analysis; M.Z. wrote the manuscript; and Y.Q.W. and S.X. revised the manuscript. All of the authors read and approved the final manuscript.

## References

Abeysinghe JK, Lam K-M, Ng DW-K. 2019. Differential regulation and interaction of homoeologous WRKY18 and WRKY40 in Arabidopsis allotetraploids and biotic stress responses. The Plant Journal 97, 352–367.

Birkenbihl RP, Kracher B, Roccaro M, Somssich IE. 2017. Induced Genome-Wide Binding of Three Arabidopsis WRKY Transcription Factors during Early MAMP-Triggered Immunity. The Plant Cell 29, 20–38.

Chen C, Chen Z. 2002. Potentiation of developmentally regulated plant defense response by AtWRKY18, a pathogen-induced Arabidopsis transcription factor. Plant physiology 129, 706–716.

Dang F, Wang Y, She J, et al.. 2014. Overexpression of CaWRKY27, a subgroup IIe WRKY transcription factor of *Capsicum annuum*, positively regulates tobacco resistance to *Ralstonia solanacearum* infection. Physiologia Plantarum 150, 397–411.

Eulgem T, Somssich IE. 2007. Networks of WRKY transcription factors in defense signaling. Current opinion in plant biology 10, 366–371.

Gadoury DM, Cadle-Davidson L, Wilcox WF, Dry IB, Seem RC, Milgroom MG. 2012. Grapevine powdery mildew (*Erysiphe necator*): a fascinating system for the study of the biology, ecology and epidemiology of an obligate biotroph. Molecular Plant Pathology 13, 1–16.

Gao YR, Han YT, Zhao FL, et al.. 2016. Identification and utilization of a new *Erysiphe necator* isolate NAFU1 to quickly evaluate powdery mildew resistance in wild Chinese grapevine species using detached leaves. Plant Physiology and Biochemistry 98, 12–24.

Guo C, Guo R, Xu X, et al.. 2014. Evolution and expression analysis of the grape (*Vitis vinifera* L.) WRKY gene family. Journal of experimental botany 65, 1513–1528.

Hu Y, Gao Y-R, Yang L-S, Wang W, Wang Y-J, Wen Y-Q. 2019. The cytological basis of powdery mildew resistance in wild Chinese *Vitis* species. Plant Physiology and Biochemistry 144, 244–253.

Hu Y, Li Y, Hou F, et al.. 2018. Ectopic expression of Arabidopsis broad-spectrum resistance gene *RPW8.2* improves the resistance to powdery mildew in grapevine (*Vitis vinifera*). Plant Science 267, 20–31.

Jaillon O, Aury J-M, Noel B, et al.. 2007. The grapevine genome sequence suggests ancestral hexaploidization in major angiosperm phyla. Nature 449, 463–467.

Jiang J, Ma S, Ye N, Jiang M, Cao J, Zhang J. 2017. WRKY transcription factors in plant responses to stresses. Journal of integrative plant biology 59, 86–101.

Liu W, Xie X, Ma X, Li J, Chen J, Liu Y-G. 2015. DSDecode: A Web-Based Tool for Decoding of Sequencing Chromatograms for Genotyping of Targeted Mutations. Molecular Plant 8, 1431–1433.

Liu X, Zhou X, Li D, Hong B, Gao J, Zhang Z. 2022. Rose WRKY13 promotes disease protection to *Botrytis* by enhancing cytokinin content and reducing abscisic acid signaling. Plant physiology 191, 679–693.

Ma T, Chen S, Liu J, et al.. 2021. *Plasmopara viticola* effector PvRXLR111 stabilizes VvWRKY40 to promote virulence. Molecular Plant Pathology 22, 231–242.

Pandey SP, Roccaro M, Schön M, Logemann E, Somssich IE. 2010. Transcriptional reprogramming regulated by WRKY18 and WRKY40 facilitates powdery mildew infection of Arabidopsis. The Plant Journal 64, 912–923.

Raffeiner M, Üstün S, Guerra T, et al.. 2022. The *Xanthomonas* type-III effector XopS stabilizes *Ca*WRKY40a to regulate defense responses and stomatal immunity in pepper (*Capsicum annuum*). The Plant Cell 34, 1684–1708.

Rushton PJ, Somssich IE, Ringler P, Shen QJ. 2010. WRKY transcription factors. Trends in Plant Science 15, 247–258.

Shen Q-H, Saijo Y, Mauch S, et al.. 2007. Nuclear activity of MLA immune receptors links isolate-specific and basal disease-resistance responses. Science 315, 1098–1103.

Tsuda K, Somssich IE. 2015. Transcriptional networks in plant immunity. The New phytologist 206, 932–947.

Wan D-Y, Guo Y, Cheng Y, et al.. 2020. CRISPR/Cas9-mediated mutagenesis of *VvMLO3* results in enhanced resistance to powdery mildew in grapevine (*Vitis vinifera*). Horticulture Research 7.

Wan R, Guo C, Hou X, et al.. 2021. Comparative transcriptomic analysis highlights contrasting levels of resistance of *Vitis vinifera* and *Vitis amurensis* to *Botrytis cinerea*. Horticulture Research 8, 103.

Wang Y, Cui X, Yang B, et al.. 2020. WRKY55 transcription factor positively regulates leaf senescence and the defense response by modulating the transcription of genes implicated in the biosynthesis of reactive oxygen species and salicylic acid in Arabidopsis. Development (Cambridge, England) 147 (16), dev189647.

Xing H-L, Dong L, Wang Z-P, et al.. 2014. A CRISPR/Cas9 toolkit for multiplex genome editing in plants. BMC Plant Biology 14, 327.

Xu X, Chen C, Fan B, Chen Z. 2006. Physical and functional interactions between pathogen-induced Arabidopsis WRKY18, WRKY40, and WRKY60 transcription factors. The Plant Cell 18, 1310–1326.

Zhao F-L, Li Y-J, Hu Y, et al.. 2016. A highly efficient grapevine mesophyll protoplast system for transient gene expression and the study of disease resistance proteins. Plant Cell, Tissue and Organ Culture (PCTOC) 125, 43–57.

